# Insane in the vembrane: filtering and transforming VCF/BCF files

**DOI:** 10.1101/2022.08.17.504106

**Authors:** Till Hartmann, Christopher Schröder, Elias Kuthe, David Lähnemann, Johannes Köster

**Affiliations:** Algorithms for reproducible bioinformatics, Genome Informatics, Institute of Human Genetics, University Hospital Essen, University of Duisburg-Essen, Essen, Germany; Genome Informatics, Institute of Human Genetics, University of Duisburg-Essen, University Hospital Essen, Essen, Germany; Computer Science XI, TU Dortmund University, Dortmund, Germany; Department of Medical Oncology, West German Cancer Center, University Hospital Essen, University Duisburg-Essen, Germany; Department of Medical Oncology, Dana-Farber Cancer Institute, Harvard Medical School, Boston, USA

## Abstract

Data from sequencing of DNA or RNA samples is routinely scanned for variation. Such variation data is stored in the standardized VCF/BCF format with additional annotations. Analyses of variants usually involve steps where filters are applied to narrow down the list of candidates for further analysis. A number of tools for this task exist, differing in functionality, speed, syntax and supported annotations. Thus, users have to switch between tools depending on the filtering task, and have to adapt to the respective filtering syntax. We present vembrane as a command line VCF/BCF filtering tool that consolidates and extends the filtering functionality of previous software to meet any imaginable filtering use case. To this end, vembrane exposes the VCF/BCF file type specification and its inofficial extensions by the annotation tools *VEP* and *SnpEff* as Python data structures. `vembrane filter` enables filtration by arbitrary Python expressions over (combinations of) annotations, requiring only basic knowledge of the Python programming language. `vembrane table` allows users to generate tables from subsets of annotations or functions thereof. Finally, it is fast, thanks to pysam, a Python wrapper around htslib, and by relying on Python’s lazy evaluation.

**Availability and Implementation:** Source code and installation instructions are available at github.com/vembrane/vembrane, DOI: 10.5281/zen-odo.7003981.

## 1 Introduction

Identifying variation from DNA or RNA sequencing data and determining its effect on phenotypes is at the heart of a wide range of biological and medical research efforts. Initial bioinformatics processing of such sequencing data routinely records thousands to millions of individual differences between one or more biological samples and their reference genome. These variants are annotated with data properties and known or predicted phenotypic effects and are usually stored in the Variant Call Format (VCF) or its binary equivalent (BCF) [Danecek et al., 2011]. This annotation information can then be used to filter down to a set of interesting candidate variants, for example those known to be drug targets in a specific disease.

Here, we present *vembrane*, a new filtering tool for all versions of the VCF and BCF formats. *vembrane* consolidates and extends the functionality of previously available tools and uses standard Python syntax, while achieving very good processing speed. The direct use of Python syntax enables flexible and powerful expressions (Fig. 1) and obviates the need to adapt to a new syntax for users already familiar with Python. It supports both *SnpEff* [Cingolani et al., 2012, Ruden et al., 2012] and *VEP* [McLaren et al., 2016] annotations out of the box and has an extensible design which allows easy integration of new annotation sources. To our knowledge, it is the only variant filtering tool that can handle groups of breakend events that represent structural variants. It consists of three subcommands for processing VCF records: `filter` for filtering, `table` for converting into a tabular format, and `annotate` for adding additional annotations based on genomic ranges.

**Figure 1:**
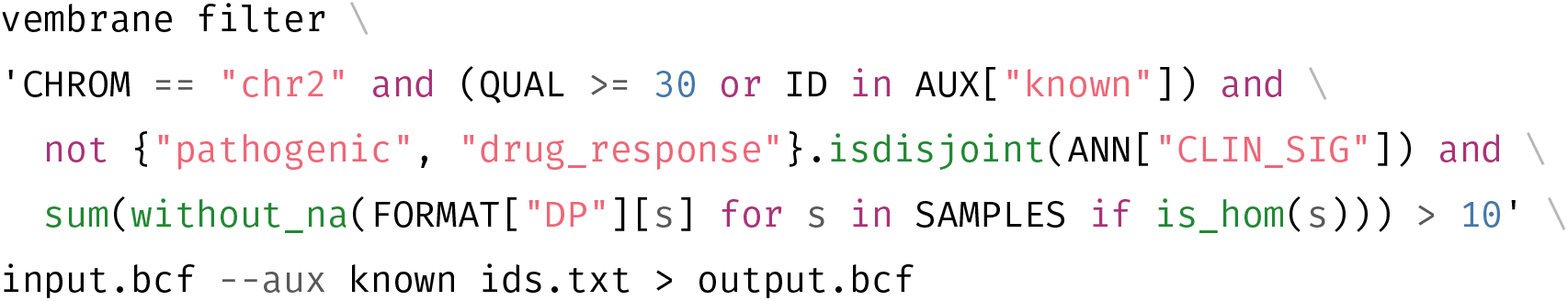
Example invocation of `vembrane filter`. The auxiliary file `ids.txt` contains one ID per line and is parsed as a set. In plain english, the filter expression translates to “keep all records from chromosome 2 where the quality is at least 30 or the ID is in the set of known IDs, and where at least ‘pathogenic’ or ‘drug_response’ is part of the clinical significance annotations, and where the sum of read depths across all samples that report a homozygous genotype is at least 10”.

## 2 Methods

### 2.1 Implementation

*vembrane* uses ordinary Python expressions with lazy evaluation. It provides all fields defined in the VCF specification as local variables, namely `CHROM`, `POS`, `ID`, `REF`, `ALT`, `QUAL`, `FILTER`, `INFO` and `FORMAT`. The entries in the `INFO` dictionary are typed according to the VCF file’s header. For annotated files, the annotation is (by default) available via the `ANN` dictionary. Annotations from *SnpEff* and *VEP* have custom parsers in vembrane, making them easier and safer to use in filter expressions. An overview of types used for these annotations is given at https://github.com/vembrane/vembrane/blob/main/docs/ann_types.md. When filtering a VCF record’s `ANN` annotation fields, vembrane by default discards `ANN` entries that do not match the expression and only discards the whole record if there are no `ANN` entries left.

For input and output, *vembrane* uses pysam as an interface to htslib [Bonfield et al., 2021]. This means *vembrane* can handle any type of VCF or BCF file, but comes with any limitation that pysam might have.

Records with multiple alternative alleles may have completely unrelated annotations. This both complicates filter expressions and interpretation of variants. Thus, *vembrane* only accepts files whose records have been split such that each alternative allele has its own record. This can for example be achieved by normalising the input with `bcftools norm -N -m-any`; for consistent results this is best done before annotation.

To our knowledge, *vembrane* is the first tool to explicitly handle breakend variants (BNDs): Breakends are a way of expressing strand breaks and their rejoining to other positions on the same reference genome. They can be used to encode structural variants by grouping two or more breakend records into a joint structural variant *event*. As variant files are usually sorted by chromosomal position, breakend records from the same event can occur in distant parts of the file. Thus, even if the event it belongs to is known for each breakend at the time of reading it, the total number of breakends (and all associated annotations) for a specific event remains unknown until reaching the end of the file. As we always want to generate valid VCF files, we need to ensure that each event is removed or kept *as a whole*, which requires an additional step for handling BNDs. For performance reasons, we yield non-BND variants instantly during iteration and defer processing of BND events until suffiicient information is available – this is the case as soon as at least one BND of an event passes the filter expression. Since this behavior does not preserve the order of input variants, it can be disabled with `--preserve-order`. This option enforces a 2-pass approach: a first pass which collects all BNDs (and skips all non-BND records), so that all groups of BNDs are known in advance for the second pass which then handles all records in order.

### 2.2 Comparison to other tools

There are a number of tools available for filtering VCF records based on conditional expressions on one or multiple fields of the VCF format. However, they vary greatly in the scope of their functionality (Table 1). For example, *SnpEff* and *VEP* annotation suites have their own filtering tools, *SnpSift* and *filter_vep*, which are tailored towards the respective annotations. Both use custom syntax, special handling of their respective annotations, and neither supports the BCF format. Additionally, *filter_vep* is several orders of magnitude slower than the other tools (suppl. Fig. S.1). The *bcftools* suite also developed its own expression syntax and supports *VEP* annotations by explicitly activating a dedicated plugin. *bio-vcf* [Garrison et al., 2021] defines its own domain specific language for processing VCF files, is multi-threaded by default, but has neither BCF support nor built-in support for annotations. *slivar* [Pedersen et al., 2021] is geared more towards trio/pedigree filter scenarios, but has some support for specific parts of *SnpEff, VEP* and *bcftools* annotations such as `Consequence`. The only other tool that does not define its own syntax is *VcfFilterJdk* [Lindenbaum and Redon, 2018], which uses Java expressions for filtering and in principle supports both *VEP* and *SnpEff* (EFF only) annotations. However, at the time of writing, it did not support VCF v4.3. A detailed comparison of specific syntactic capabilities of the different tools, as well as a performance benchmark, can be found in the supplement.

**Table 1:** Overview of different tools and their properties. See supplement for detailed benchmarking. ^a^via +split-vep plugin, ^b^additionally “*custom*” because some scenarios require complex CLI option combinations, ^c^special handling of impact annotations from *bcftools, VEP* or *SnpEff*, ^d^EFF only, ^e^VCF *<* v4.3 only, ^f^manually estimated performance, since it is not included in the benchmark due to incompatible VCF version support and lack of conda packages.

| tool | syntax | annotation | I/O formats | breakends | speed |
| --- | --- | --- | --- | --- | --- |
| *bcftools* | custom | *VEP*<sup>a</sup> | VCF, BCF | no | +++ |
| *bio-vcf* | custom/ruby<sup>b</sup> | – | VCF | no | – |
| *filter_vep* | custom | *VEP* | VCF | no | --- |
| *slivar* | js/custom<sup>b</sup> | custom<sup>c</sup> | VCF, BCF | no | – |
| *SnpSift* | custom | *SnpEff* | VCF | no | + |
| *VcfFilterJdk* | Java | *VEP*, *SnpEff*<sup>d</sup> | VCF, BCF<sup>e</sup> | no | ○<sup>f</sup> |
| *vembrane* | Python | *VEP*, *SnpEff* | VCF, BCF | yes | ++ |

## 3 Summary

*vembrane* is a new software for effiicient filtering of variation data in the standardized VCF and BCF formats. It combines the capabilities of existing tools and should work as a replacement to any of them. Thus, users will not have to remember which tool can achieve what, but should be able to perform any filtering task with *vembrane*. Further, *vembrane* allows for filtering via arbitrary Python expressions, meaning that Python users can compose filtering expressions without having to learn custom syntax. In addition, it extends beyond existing functionality in other tools by providing support for breakends. Finally, it also allows formatting VCF files into tables and has basic support for annotating records itself.

## Supporting information

Benchmark

## 4 Acknowledgements

We sincerely thank Marcel Bargull, Jan Forster, Felix Mölder and Sven Rahmann for their contributions.

