## Supplementary material for "Insane in the vembrane: filtering and transforming VCF/BCF files": Benchmark

In order to compare different VCF filtering tools with respect to processing times, we ran them on VCF files with different numbers of samples. These were generated by annotating the GIAB VCF files[1] (restricted to chromosome 1) of the samples `HG001`, `HG002`, `HG003` and `HG004` with both *SnpEff* and *VEP*, and creating VCF files with all possible multi-sample combinations of 1, 2, 3 or 4 samples. These were then filtered with tools that adhere to the specifications of VCF in version 4.3 and BCF in version 2.2[2], the versions htslib and bcftools use at the time of writing. This decision rules out *VcfFilterJdk* since htsjdk only supports earlier versions of the VCF/BCF specifications and hence produces incompatible files.

[1] https://www.nist.gov/programs-projects/genome-bottle

[2] https://samtools.github.io/hts-specs/VCFv4.3.pdf

We selected a range of different filter expressions, varying in field accesses and general expression complexity, see Table S.1 for details. We then benchmarked each tool on the annotated files with 10 repeats. Results are shown in Fig. S.1.

To make sure the same records were kept, we computed md5sums on the filtered and sorted VCF files while restricting fields to `CHROM`, `POS`, `REF`, `ALT` and `QUAL`. This restriction is necessary because:

- *bcftools* with `+split-vep` will add `INFO` fields for every field parsed from the `ANN` annotation by default
- *SnpSift* keeps all annotations if at least one of them matches (*vembrane* can mirror this behaviour with `--keep-unmatched`)
- *slivar* adds `impactful`, `genic` and `highest_impact_order` `INFO` fields

To reproduce the results, the snakemake workflow used for benchmarking is available at `github.com/vembrane/vembrane-benchmark` (`10.5281/zenodo.6979842`).

The benchmark (as shown in Fig. S.1) was run on a machine with an Intel(R) Xeon(R) Gold 6152 Processor (88 cores) with 768GiB RAM and 160TB of harddisk space managed in LVM groups. At any given time, a maximum of 4 jobs were run in parallel to limit I/O load.

| scenario name | tool | expression |
|---|---|---|
| filter_all | vembrane | False |
| | SnpSift | false |
| | slivar | --info 'false' |
| | filter_vep | 0 |
| | bio-vcf | false |
| | bcftools | -e "" |
| filter_none | vembrane | True |
| | SnpSift | true |
| | slivar | --info 'true' |
| | filter_vep | not 0 |
| | bio-vcf | true |
| | bcftools | -i "" |
| at_least_2_platforms | vembrane | INFO["platforms"] >= 2 |
| | SnpSift | platforms >= 2 |
| | slivar | --info 'INFO.platforms >= 2' |
| | filter_vep | platforms >= 2 |
| | bio-vcf | rec.info.platforms >= 2 |
| | bcftools | -i "INFO/platforms >= 2" |
| format_dp | vembrane | any(FORMAT["DP"][s] > 1250 for s in SAMPLES) |
| | SnpSift | GEN[*].DP > 1250 |
| | slivar | --alias resources/empty_alias.txt --pass-only --sample-expr ':sample.DP > 1250' |
| | filter_vep | cannot access FORMAT |
| | bio-vcf | --sfilter defaults to conjunctions ("all"), not disjunctions ("any") |
| | bcftools | -i "FORMAT/DP > 1250" |
| impact_high | vembrane | ANN["Annotation_Impact"] == "HIGH" |
| | SnpSift | ANN[*].IMPACT has 'HIGH' |
| | slivar | SLIVAR_IMPACTFUL_ORDER=slivar-impactfulness-order.txt slivar expr --info "INFO.impactful" |
| | filter_vep | ignores SnpEff (or any non-VEP) annotation without raising an error |
| | bio-vcf | No built-in support for annotations |
| | bcftools | No built-in support for *SnpEff* annotations |
| uncertain | vembrane | "uncertain_significance" in ANN["CLIN_SIG"] or not (ID and ID.startswith("rs")) |
| | SnpSift | cannot access *VEP* annotations |
| | slivar | No built-in support for annotations apart from "Consequence" |
| | filter_vep | CLIN_SIG is uncertain_significance or not (ID and ID matches "^rs" ) |
| | bio-vcf | No built-in support for annotations |
| | bcftools | +split-vep --annotation "ANN" -c CLIN_SIG -i "INFO/CLIN_SIG[*] == 'uncertain_significance' \| (ID \!~ '^rs')" |

Table S.1: Expressions used for benchmarking. `impact_high` makes use of *SnpEff* annotations, `uncertain` makes use of VEP annotations, all other expressions only use default VCF fields and/or `INFO` and `FORMAT` fields defined in the header. For some tools it is necessary to specify the commandline options as well, e. g. for *bcftools* the interpretation of the expression changes: `-i` *includes*, `-e` *excludes* variants matching the expression.

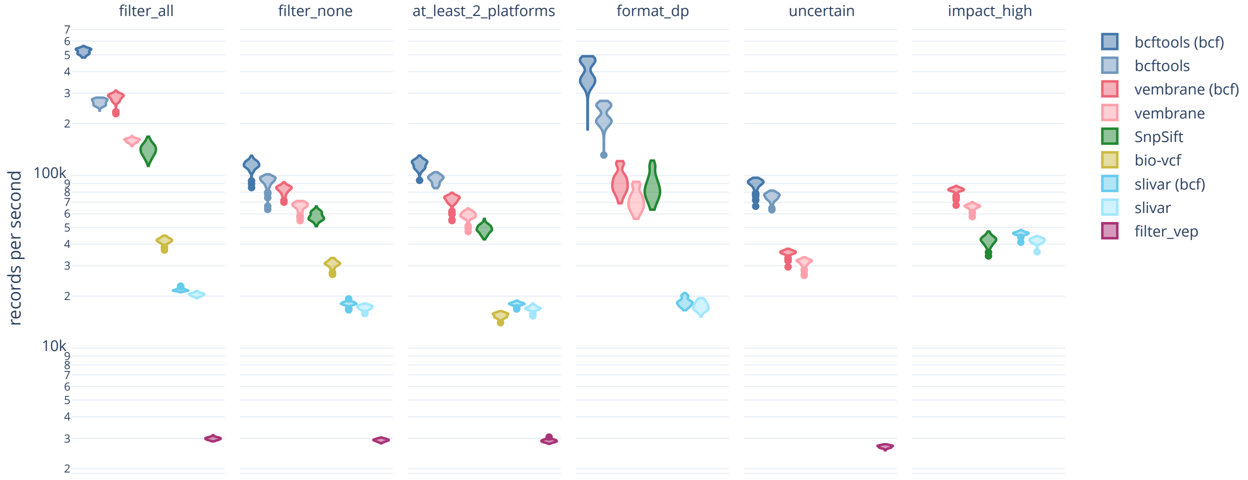


Figure S.1: A benchmark comparing *vembrane*, *bcftools*, *SnpSift*, *filter_vep*, *bio-vcf* and *slivar*. The $y$-axis is in records per seconds, i. e. higher is better. Runs with BCF input are listed separately for tools that support this. Each column corresponds to a different filter expression as described in Table S.1. Note the logarithmic scale on the $y$-axis.

### *Pitfalls*

Different VCF filtering tools have different use-cases and focuses and provide different levels of convenience abstractions and conventions. This becomes especially apparent in the following categories, each illustrated with examples:

#### *Multi-allelic records*

The VCF file format allows multiple alternate alleles per record (i. e. per line in the textual representation). Because working with multiple alternate alleles quickly gets verbose, tools often use implicit conjunctions or disjunctions.

Consider the following (incomplete) example VCF file with two alternate alleles and read depth information in the info field `DP`:

```
##INFO=<ID=DP,NUMBER=A,Type=Integer,Description="Depth">
#CHROM  POS REF ALT INFO
ctg1    42  A   C,G DP=0,50
```

Now, with the *bcftools* expression `INFO/DP > 0` the record is *kept*, because `INFO/DP > 0` is defined implicitly as `INFO/DP[*] > 0`. The square brackets denote array subscript and the asterisk denotes ”any element”. This translates to ”at least one alternate allele in the record must have read depth greater than zero”.
However, with the *SnpSift* expression `DP > 0` the record is *discarded*, because unlike *bcftools*, *SnpSift* implicitly assumes `DP[?] > 0`. Here, the question mark implies ”all elements” must match the (respective part of the) expression. This translates to ”all alternate alleles in the record must have read depth greater than zero”.

We argue that multi-allelic records should be split into multiple records with only one alternate allele each, and annotation of such records should only happen after splitting, since annotations may be completely unrelated between different alternate alleles. This also eliminates one source of differences between implicit behaviour of VCF filtering tools.

#### *Conventions*

Usually certain syntactical elements imply *conventional* semantics. For example, `=` and `==` are often either used as variable assignment and comparison operator respectively or both interpreted as comparison operators. Similarly, logical operators such as `&` or `&&` (and, conjunction) or `|` and `||` (or, disjunction) can often be used interchangeably[3] *without changing semantics.*
In *bcftools* syntax, both `&` and `&&` are logic operators denoting a conjunction: both `FMT/DP > 0 & FMT/GQ > 10` and `FMT/DP > 0 && FMT/GQ > 10` are valid expressions. The former translates to "read depth `> 0` and genotype quality `> 10` must be fulfilled for the same sample". The latter, however, translates to "read depth `> 0` in any sample and genotype quality `> 10` in any sample". When written as *vembrane* expressions, the difference between the two can be seen easily:

```
 &: any((FORMAT["DP"][s] > 0 and FORMAT["GQ"][s] > 10) for s in SAMPLES)
&&: any(FORMAT["DP"][s] > 0 for s in SAMPLES) and any(FORMAT["GQ"][s] >
10 for s in SAMPLES)
```

Therefore, `&` and `&&` cannot be used interchangeably in *bcftools* because they have different semantics.

On a related note, using `&` or `&&` in *filter_vep* expressions does not error (and simply yields every record, unfiltered), even though the only valid conjunction is expressed using `and`.

[3] In many languages, singular `&` and `|` denote bitwise *and* and *or*, while double `&&` and `||` denote logical *and* and *or*.

#### *Portability*

In contrast to *bcftools*, *vembrane* does *not* automatically collapse/elevate `INFO` vs. `FORMAT` fields if they are unambiguous: For example, in *bcftools* the expression `platforms > 3` is valid as long as there is only one field named `platforms`; as soon as there is ambiguity, the full path has to be specified, so either `INFO/platforms > 3` or `FORMAT/platforms > 3`. This is a portability issue, as the same expression used in a different context may error or silently behave differently than intended. Here, and in general, *vembrane* is stricter than other tools (and has fewer special cases). As a consequence, the expressions are more portable and their interpretation less ambiguous.

### *Expressions*

#### *Syntax*

A formal introduction and description of python's syntax can be found at `https://docs.python.org/3/reference/expressions.html`.

In Table S.2 we present a showcase of various vembrane/VCF specific example expressions and their interpretations.

| expression | interpretation |
| --- | --- |
| `CHROM == "chr2"` | select variants on chromosome 2 |
| `QUAL >= 30 or ID in AUX["known_ids"]` | require either quality at least 30 or ID contained in auxiliary set |
| `any(e in ANN["CLIN_SIG"] for e in ("pathogenic", "drug_response"))` | at least ”pathogenic” or ”drug_response” in the list of clinical significances |
| `re.search(r"(up|down)stream", ANN["Consequence"])` | consequence should contain either ”upstream” or ”downstream” |
| `all(FORMAT["AF"][s] >= 0.5 for s in SAMPLES)` | allele frequency at least 0.5 for all samples |
| `all(v >= 0.5 for v in FORMAT["AF"])` | same as above |
| `sum(without_na(FORMAT["DP"][s] for s in SAMPLES)) > 10` | sum of read depth across samples with read depth information at least 10 |
| `sum(without_na(FORMAT["DP"][s] for s in SAMPLES if is_hom(s))) > 10` | same as above but additionally restricts to samples that report a homozygous genotype |

Table S.2: Some example *vembrane* expressions with their corresponding interpretations.

#### *Semantic equivalency*

Since *vembrane* relies on python for evaluation, it is possible to express the same filter as semantically equivalent but syntactically different expressions, for example:

```
any(entry in ANN["CLIN_SIG"] for entry in ("pathogenic",
↪  "likely_pathogenic", "drug_response"))
```

or equivalently

```
not {"pathogenic", "likely_pathogenic",
↪  "drug_response"}.isdisjoint(ANN["CLIN_SIG"])
```

or

```
"pathogenic" in ANN["CLIN_SIG"] \
↪  or "likely_pathogenic" in ANN["CLIN_SIG"] \
↪  or "drug_response" in ANN["CLIN_SIG"]
```

While these expressions are semantically equivalent, the python interpreter will not always perform the same operations; this may result in differences in performance. We recommend using expressions that are easy to understand by humans while still remaining concise.

#### *Data availability statement*

The URLs to the VCF files used for benchmarking are listed in Table S.3.

The benchmark workflow is available at `github.com/vembrane/vembrane-benchmark` (`10.5281/zenodo.6979842`).

| Sample | URL |
|---|---|
| HG001 | ftp://ftp-trace.ncbi.nlm.nih.gov/giab/ftp/release/NA12878_HG001/latest/GRCh38/HG001_GRCh38_GIAB_highconf_CG-IllFB-IllGATKHC-Ion-10X-SOLID_CHROM1-X_v.3.3.2_highconf_PGandRTGphasetransfer.vcf.gz |
| HG002 | ftp://ftp-trace.ncbi.nlm.nih.gov/giab/ftp/release/AshkenazimTrio/HG002_NA24385_son/latest/GRCh38/HG002_GRCh38_1_22_v4.1_draft_benchmark.vcf.gz |
| HG003 | ftp://ftp-trace.ncbi.nlm.nih.gov/giab/ftp/release/AshkenazimTrio/HG003_NA24149_father/latest/GRCh38/HG003_GRCh38_GIAB_highconf_CG-Illfb-IllsentieonHC-Ion-10XsentieonHC_CHROM1-22_v.3.3.2_highconf.vcf.gz |
| HG004 | ftp://ftp-trace.ncbi.nlm.nih.gov/giab/ftp/release/AshkenazimTrio/HG004_NA24143_mother/latest/GRCh38/HG004_GRCh38_GIAB_highconf_CG-Illfb-IllsentieonHC-Ion-10XsentieonHC_CHROM1-22_v.3.3.2_highconf.vcf.gz |

Table S.3: FTP URLs of VCF files for GIAB samples `HG001`, `HG002`, `HG003` and `HG004`

### *References*

Till Hartmann and David Lähnemann. vembrane/vembrane-benchmark: v1.1.0, Jul 2022.
